# A portable and cost-effective microfluidic system for massively parallel single-cell transcriptome profiling

**DOI:** 10.1101/818450

**Authors:** Chuanyu Liu, Tao Wu, Fei Fan, Ya Liu, Liang Wu, Michael Junkin, Zhifeng Wang, Yeya Yu, Weimao Wang, Wenbo Wei, Yue Yuan, Mingyue Wang, Mengnan Cheng, Xiaoyu Wei, Jiangshan Xu, Quan Shi, Shiping Liu, Ao Chen, Ou Wang, Ming Ni, Wenwei Zhang, Zhouchun Shang, Yiwei Lai, Pengcheng Guo, Carl Ward, Giacomo Volpe, Lei Wang, Huan Zheng, Yang Liu, Brock A. Peters, Jody Beecher, Yongwei Zhang, Miguel A. Esteban, Yong Hou, Xun Xu, I-Jane Chen, Longqi Liu

**Affiliations:** BGI-Shenzhen, Shenzhen 518083, China; China National GeneBank, BGI-Shenzhen, Shenzhen 518120, China; BGI Education Center, University of Chinese Academy of Sciences, Shenzhen 518083, China; Complete Genomics Inc., 2904 Orchard Parkway, San Jose, California 95134, USA; BGI College, Zhengzhou University, Zhengzhou 450000, China; Laboratory of Genomics and Molecular Biomedicine, Department of Biology, University of Copenhagen, Copenhagen 2200, Denmark; MGI, BGI-Shenzhen, Shenzhen 518083, China; Laboratory of RNA, Chromatin, and Human Disease, Guangzhou Institutes of Biomedicine and Health, Chinese Academy of Science, Guangzhou 510530, China; College of Veterinary Medicine, Jilin University, Changchun 130062, China; Institute for Stem cell and Regeneration, Chinese Academy of Sciences, Beijing 100101, China

## Abstract

Single-cell technologies are becoming increasingly widespread and have been revolutionizing our understanding of cell identity, state, diversity and function. However, current platforms can be slow to apply to large-scale studies and resource-limited clinical arenas due to a variety of reasons including cost, infrastructure, sample quality and requirements. Here we report DNBelab C4 (C4), a negative pressure orchestrated, portable and cost-effective device that enables high-throughput single-cell transcriptional profiling. C4 system can efficiently allow discrimination of species-specific cells at high resolution and dissect tissue heterogeneity in different organs, such as murine lung and cerebral cortex. Finally, we show that the C4 system is comparable to existing platforms but has huge benefits in cost and portability and, as such, it will be of great interest for the wider scientific community.

## Introduction

The emergence of single-cell sequencing technologies has offered great promises in the interrogation of tissue heterogeneity that could not previously be resolved from measuring the average gene expression of a bulk cell population [1]. During the past 10 years, single-cell genomics technologies have evolved rapidly at scale and power, enabling the profiling of thousands to tens of thousands single-cells per experiment [2]. These efforts have enabled a comprehensive characterization of tissue heterogeneity, developmental trajectories, cellular reprogramming and human diseases at an unprecedented resolution.

Current high-throughput single-cell technologies apply droplet and micro-well based methods, which enable partition and barcoding of single cells inside nano-liter reactors. Micro-particles coated with barcoded oligos are employed to capture mRNA or DNA molecules from each cell [3-5]. These methods, in comparison to low-throughput plate-based methods, are robust in cell type classification since profiling higher number of cells can reduce the negative impact from technical and intrinsic noise [6]. However, the effectiveness of these methods is limited by lengthy approaches and by the use of expensive commercial instruments and reagents. Specifically, clinical samples require to be transported to different laboratories for library preparation because current microfluidic single-cell platforms are not portable. To address these short-comings, we have developed DNBelab C4, a hand-held microfluidic device to perform cell separation before high-throughput single-cell transcriptional profiling. C4 is a vacuum driven and user-friendly droplet system which offers a cost-effective and portable single-cell technology for both basic research and clinical purposes. C4 is also an extensible platform that can be further implemented for a variety of high-throughput omics technologies, such as scATAC-seq and scChIP-seq. Here we show that the C4 system is comparable to existing platforms but has benefits in cost and portability.

## Results

### A hand-held and cost-effective microfluidic system for single-cell profiling

We have previously developed a hand-held, power-free microfluidic device that can stably generate mono-dispersion sub-nanoliter size droplets [7]. To further apply this system to single-cell profiling, we have optimized both the structure of microfluidic chip and parameters of negative pressure to ensure high compatibility with single-cell RNA sequecing (scRNA-seq) related reagents. To this end, we developed C4 system (Fig. 1a), which is a low cost device composed of a syringe, a microfluidic chip, and a station. C4 is a fully manual system and does not have any electronic parts. The syringe is connected to the chip via connection tubings and placed on the station. After all reagents and samples are loaded in the inlet reservoirs, the plunger of the syringe is pulled to generate negative pressure that drives the reagents flowing into the fluidic passage to form droplets. This single source negative pressure ensures the flows of all reagents occur simultaneously without any fluctuation or lag in flow rates. The air volume in the syringe before and after plunger is pulled determines the degree of vacuum, which follows ideal gas laws. The station holds the plunger in place after it is pulled so the vacuum is uniformly maintained. To evaluate the uniformity of droplets size, more than 1.5 million droplets each run were generated independently from 4 users using 14 chips; we observed an average size of 55.9 μm with a standard deviation of 2.4 μm (Fig 1b, n = 14,000), revealing high reproducibility and stability of the negative pressure-driven system in generating large numbers of droplets.

**Figure 1.**
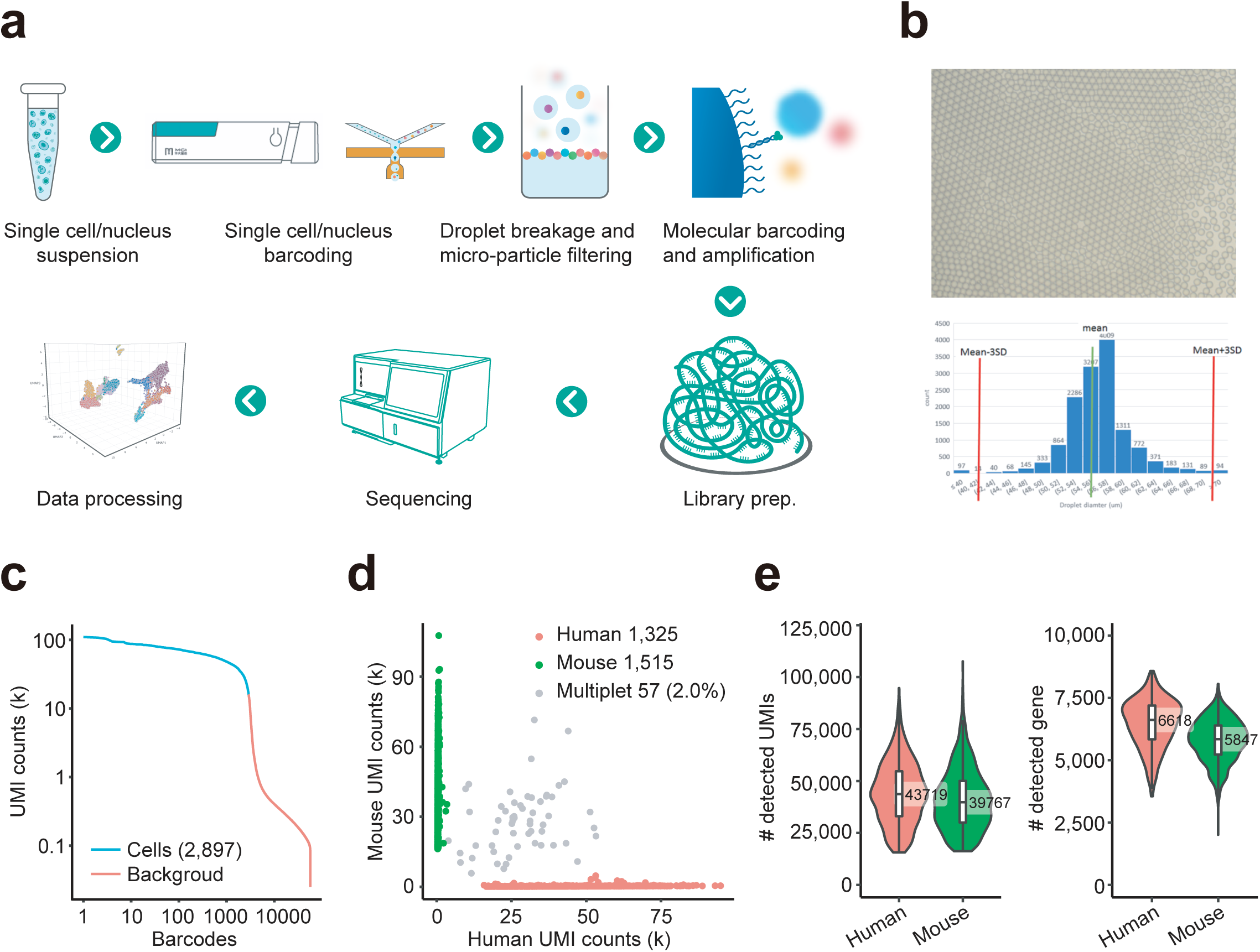
Massively parallel single-cell transcriptome profiling using C4 system. (**a**) Schematic representation of the workflow for scRNA-seq using C4 system. (**b**) Upper panel: an image of droplets generated by C4 system. Bottom panel: distribution of the size of droplets generated by different users and from different chips. (**c**) Number of UMIs captured in each cell barcode. The cell barcodes are ordered by the UMIs number. (**d**) Scatter plot of human and mouse UMI counts detected in a mixture of 293T and NIH3T3 cells. Cell barcodes containing human reads are colored in red and termed ‘Human’; cell barcodes with mouse reads are colored in green and termed ‘Mouse’; and cell barcodes with significant human reads are coloured in light grey and termed ‘Multiplet’. The percentage of multiplets detected in indicated in the panel (2%). (**e**) Left panel: number of UMIs per cell for the human and mouse cells. Right panel: number of genes per cell for the human and mouse cells.

### Massively parallel scRNA-seq using C4 system

We next developed an entire workflow for scRNA-seq using C4 system (Fig 1a). Cells, functionalized beads and lysis buffer were encapsulated into emulsion droplets, where each cell was lysed and mRNA transcripts were captured by bead-linked single-strand oligonucleotides consisting of sequencing adaptor, cell barcode, unique molecular identifier (UMI) and oligo-dT. Emulsion droplets were then transferred into a membrane filter (pore size: 8 μm), where emulsion was broken and filtered by either negative pressure or centrifugation. Reverse transcription of mRNA transcripts was conducted on pooled beads, resulting in cDNA molecules containing both cell barcode and UMIs. Subsequently, the cDNA molecules were amplified and sheared for short-read based sequencing library preparation.

To assess the technical performance of C4 system in single-cell transcriptional sequencing, we profiled cells from a mixture of 50% human (HEK293T) and 50% mouse (NIH3T3) cells at the concentration of 1,000 cells/μl. A total of 548 million quality-filtered reads were obtained and then assigned to each cell barcode and were mapped to reference genomes. 61% of the reads mapped to exonic regions (Supplementary Fig. 1a), revealing high library quality. One library contained an estimated 2,897 cells based on the distribution of the number of UMIs per barcode (Fig. 1c). A discrimination between mouse and human UMI revealed only a small percentage (roughly 2%) of cells that contain high fraction of both human and mouse reads (Fig. 1d). In addition, an average of 43,719 and 39,767 UMI-tagged transcripts (assigned to 6,618 and 5,847 genes) were detected in human and mouse cells, respectively (Fig. 1e). These data demonstrate that C4 system shows minimal collision rate and high gene capture efficiency. We performed an independent cross-species experiment and observed similar results (Supplementary Fig. 1b-d) and strong correlations between replicates (Pearson correlation = 0.998, Supplementary Fig. 2), thus strenghening the effectiveness of our approach.

**Figure 2.**
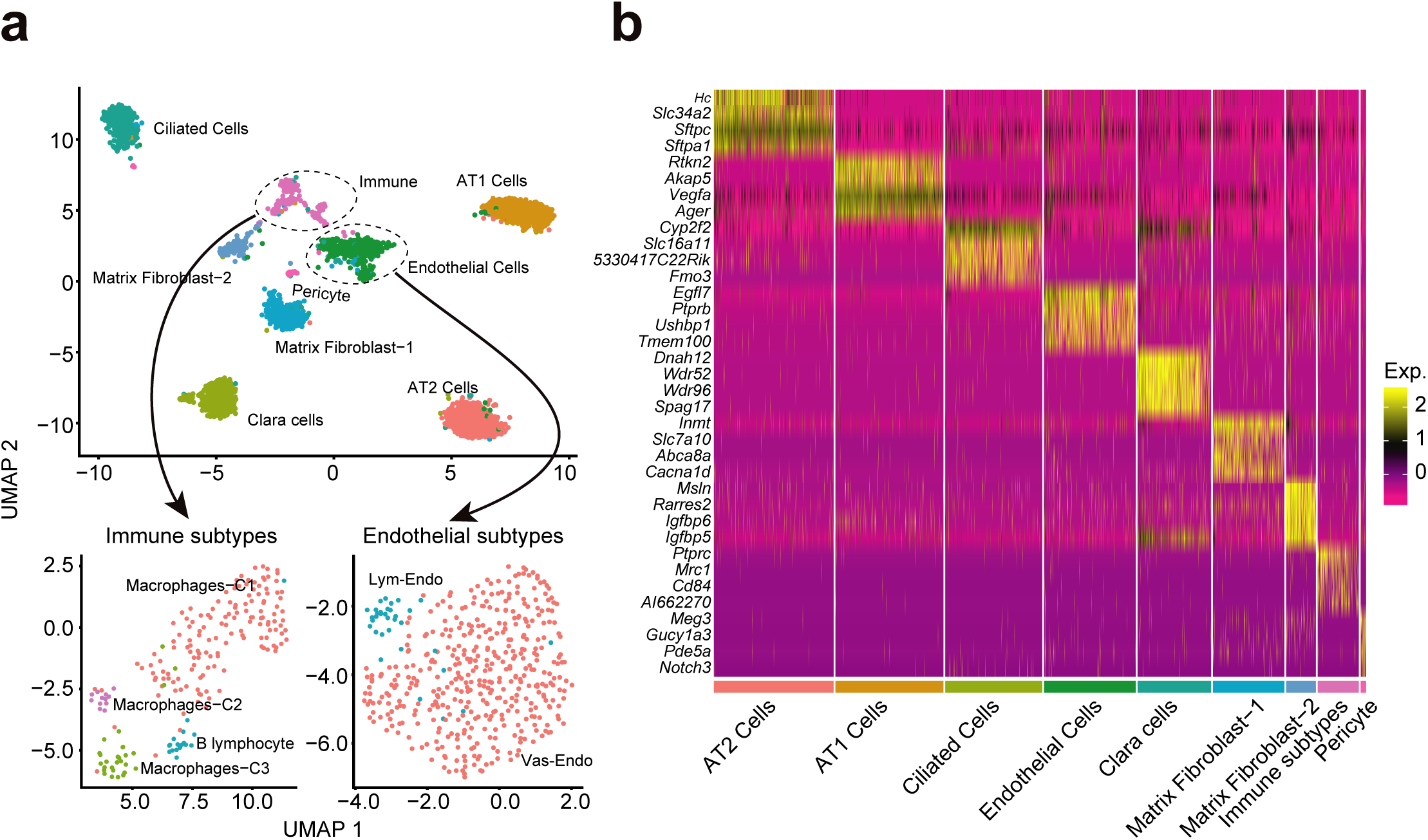
C4 can recover diverse cell types from heterogeneous samples. (**a**) Unsupervised clustering identified 13 distinct subpopulations in mouse lung, which were then assigned to cell types based on gene expression, including 4 immune subtypes and 2 endothelial subtypes. (**b**) Heatmap showing differential gene expression analysis result that specifically expressed in each subpopulation.

### Unsupervised taxonomy of cellular states

To examine the ability of C4 system to resolve cell populations in complex primary tissues, we isolated nuclei, an extensively adopted type of input sample [7-9], from mouse lung tissue to generate single-nucleus RNA sequecing (snRNA-seq) libraries. After sequencing and read alignment, we collected a total of 3,241 cells based on the UMI distribution (data not shown). Unsupervised clustering identified 9 distinct subpopulations, suggesting high heterogeneity within this tissue (Fig. 2a). Differential gene expression analysis identified genes that were specifically expressed in each subpopulation (Fig. 2b), resulting in the characterization of cell types including epithelial cells such as alveolar type 1 (AT1, expressing *Ager*) and type 2 (AT2, expressing *Sftpb*) cells, endothelial cells (*Pecam1*), fibroblast cells (*Igfbp5* and *Fn1*), and immune cells (*Ptprc*) (Fig. 2a, Supplementary Fig. 3). Interestingly, 4 sub-types of cells can be further identified within immune cells including 3 types of macrophages and one type of B lymphocytes, while endothelial cell population can also be further sub-classified into those of vascular and lymphocyte of origin, respectively (Fig. 2a). These data are consistent with previous single-cell analysis of lung tissue from newborn mice [10], suggesting that C4 system could be faithfully applied into dissecting complex cell populations at high resolution.

**Figure 3.**
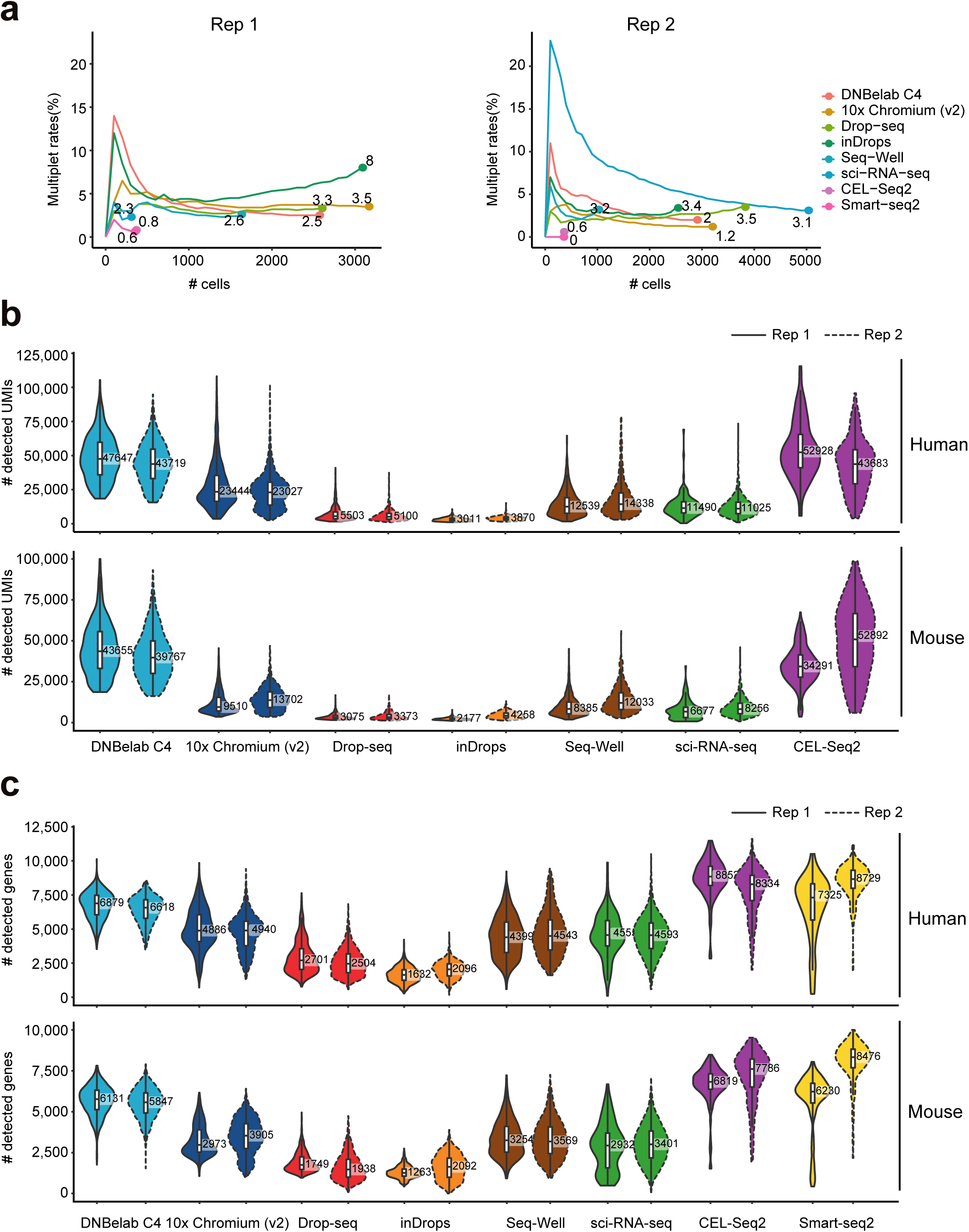
Comparative analysis of C4 and existing platforms. (**a**) Assessment of multiplet frequency. Cells were ordered based on the number of UMIs. For a given x value, the plot shows the percent of the top x cells that are multiplets. (**b**) Violin plots showing the number of UMIs per cell in each method. (**c**) Violin plot showing the number of genes per cell in each method. For (**b**) and (**c**), the top panel shows the results for human cells, and the bottom panel shows the results for mouse cells. Boxplots denote the medians (labelled on the right) and the interquartile ranges (IQRs).

### Comparative analysis of C4 and existing platforms

Two recent benchmarking studies by Ding *et al.* and Mereu *et al*. [11, 12] have systematically and comprehensively compared existing single-cell sequencing techniques, providing invaluable resource for users to make informed choices. To compare the technical performance between C4 and these single-cell platforms, we firstly analyzed the data from HEK293T/NIH3T3 mixture cells in replicates and the data generated from the same cell types in the study by Ding *et al*. [11]. We observed a multiplet rate of 2-2.5% in C4 system, which is quite comparable and acceptable in comparison with other high-throughput platforms (Fig. 3a). To evaluate the gene detection sensitivity, we first performed down-sampling analysis and found that a median number of over 6,000 genes was identified at 100K reads per cell, which outperforms other platforms (Supplementary Fig. 4). Supporting this, we further calculated the number of UMIs and genes of all single cells from each platform. As expected, the plate-based method, Smart-seq2 and CEL-Seq2 showed the highest capture sensitivity (Fig. 3b, c). Interestingly, when comparing with high-throughput methods, C4 detected over 6,000 genes in both mouse and human cell lines, which is much higher than all other platforms (Fig. 3b, c), possibly owing this to the higher number of capture oligos (theoretical number: 10^7^) on our functionalized micro-particles.

**Figure 4.**
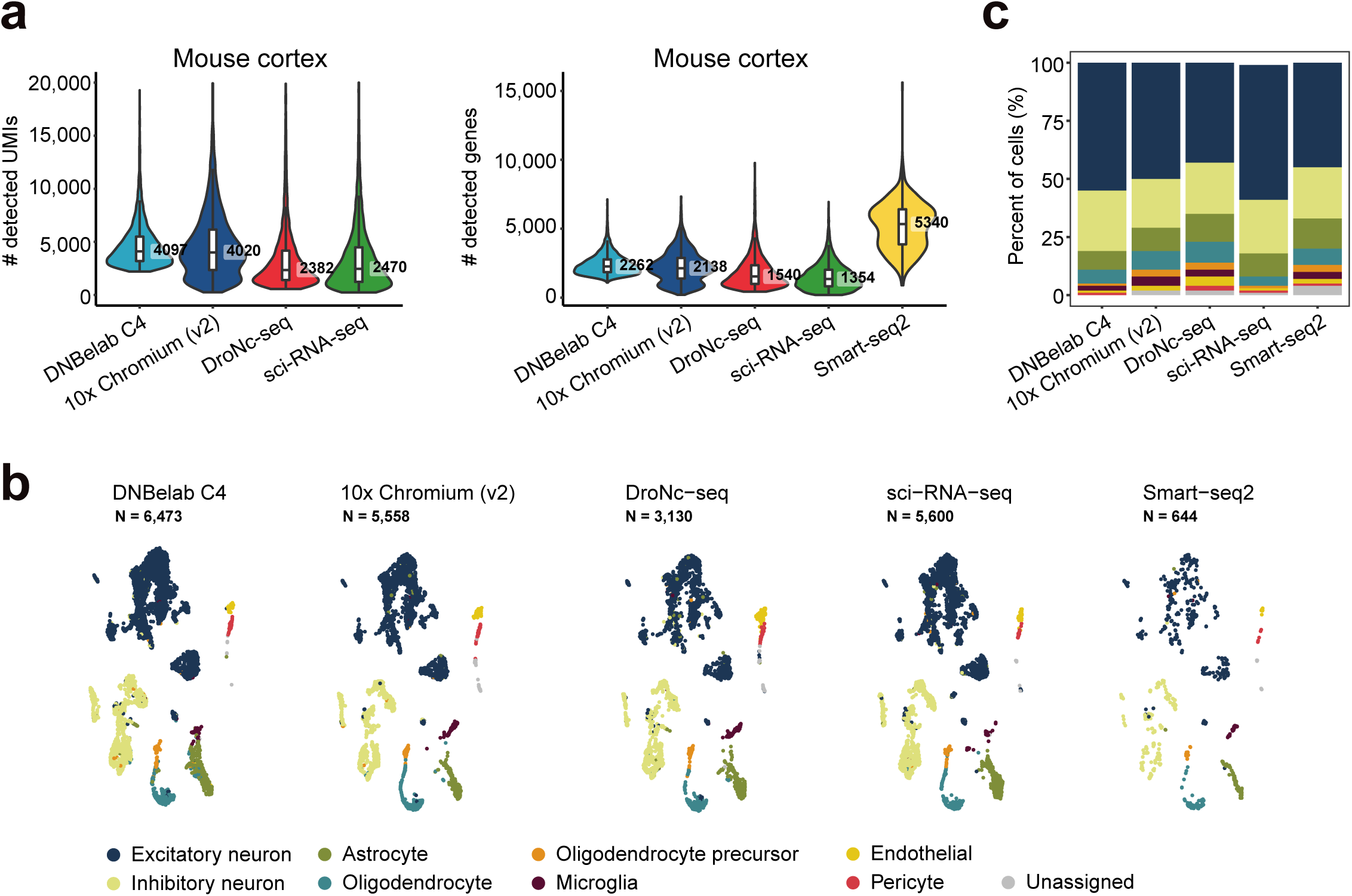
Comparative analysis of C4 and existing systems in the recovery of different cell types within mouse brain cortex. (**a**) Left panel: Violin plot showing the number of UMIs per cell from mouse cortex snRNA-seq data. Right panel: Violin plot showing the number of genes per cell in data generated by each method. (**b**) Cell clustering and cell type annotation for data generated by each method. (**c**) Proportion of the indicated cell types detected from the data by each method.

One of the key biological information obtained from scRNA-seq is the identification and recovery of different cell types from heterogeneous populations. In their report, Ding *et al*. [11] compared the ability of different platforms to resolve cellular heterogeneity in mouse cortex and human PBMCs. To further compare this facet of C4 with current platforms, we profiled a total of 6,473 single cells from the nuclei of mouse cortex, which was disassociated and extracted according to the protocol by Ding *et al*. [11]. As expected, we observed slightly higher number of genes in C4 system (Fig. 4a) than all other high-throughput platforms. We next performed unsupervised clustering for the nuclei data based on gene expression matrices, resulting in 9 subpopulations. Marker gene based annotation revealed recovery of known cell types in mouse cortex in all methods, e.g., excitatory or inhibitory neurons, astrocytes and microglia (Fig. 4b). We further calculated the percentage of each subpopulation and observed successful capture of all major cell types (Fig. 4c). C4 and other droplet-based methods can recover all cell types, whereas the combinatorial indexing based method, sci-RNA-seq, can not detect cell types such as pericytes, suggesting potential bias across different strategies. Taken together, these results strongly indicated that C4 presents high sensitivity in gene detection and comparable recovery ability with most of the current platforms.

## Discussion

In summary, we have developed a portable, affordable and user friendly microfludic system that enables scalable single-cell transcriptomics profiling. The portability feature of C4 system is critical since many samples should be freshly prepared and immediately loaded to ensure high data quality. C4 is applicable to different tissues and sample types including viable cells and nuclei, which is important since only nuclei can be isolated from many archived samples including frozen tissues or dissociated cells. In addition, we have systematically compared C4 and other 7 single cell platforms, demonstrating that C4 shows high sensitivity on mRNA molecule detection and robust ability for recovery of cellular heterogeneity in different tissues, such as cortex. Apart from good technical performance, C4 system is of extremely low-cost, making it possible for every life science lab to do single-cell studies without purchasing expensive instruments.

One limitation for current high throughput single-cell platforms (such as droplet, microwell or combinatorial indexing based methods) is that only 3’ or 5’ end exonic read can be measured on short-read based sequencing platforms. The tag information can only be used for expression quantification but does not allow transcriptomic analyses such as alternative splicing and structural variation in a high-throughput manner [1]. However, full-length cDNA library can be obtained from these platforms including C4 system. One potential approach to achieve full-length cDNA sequencing is to combine long read sequencing techniques such as long fragment read (LFR) technology [13], which has been widely used for DNA sequencing. Moreover, C4 could be an extendable platform for efficient profiling of other omics layers, such as chromatin accessibility or simultaneous profiling of multi-omics at single-cell resolution, with specific modifications to oligo on the micro-particles. Overall, C4 is a promising candidate single-cell system for high-throughput multimodal study of single cells, paving the way to a thorough assessment of cellular heterogeneity in a variety of basic research and clinical applications.

## Methods

### Cell lines and single cell suspension preparation

293T human embryonic kidney cells (ATCC) and NIH3T3 mouse embryonic fibroblast cells (ATCC) were cultured in medium containing Dulbecco’s Modified Eagle medium (Gibco) supplemented with 1X penicillin-streptomycin and 10% fetal bovine serum (Gibco).

Cells were grown to a confluence of 50-60%. Cells were treated with Trypsin-EDTA (Thermo Fisher Scientific) for 5 minutes, quenched with equal volume of complete growth medium, and spun down at 300 x g for 5 minutes. The supernatant was removed, and cells were washed twice with 1X phosphate buffered saline (PBS). Then cells were resuspended in 1X PBS containing 0.01% bovine serum albumin (BSA, BBI), passed through a 40 μm cell strainer (Falcon) and then centrifuged at 300 x g for 5 minutes. Cells were resuspended with cell resuspension buffer (MGI) at a concentration of 1,000 cells/μl. To evaluate the collision rate of C4 system, we mixed the 293T and NIH3T3 cells at a 1:1 ratio as input sample.

### Mouse tissues dissection and nuclei isolation

Wild-type C57BL/6J male mice were purchased from Guangdong Medical Lab Animal Center (Guangzhou, China). All experiments in this study were approved by the Institutional Review Board on Ethics Committee of BGI. We obtained lung and cortex tissues from 1 month old C57BL/6 male mice. The lung tissue was extracted and cut into 6 pieces (50-200 mg/piece). For cortex, we dissected the whole cortex out of the first hemisphere by a midsagittal cut and a cut made between the cerebellum and the hemisphere. The hippocampus, olfactory bulb, and all basal ganglia were dissected out as previously described [11]. The cortex and the lung slides were placed in 500 μl RNAlater (Thermo Fisher Scientific) in 1.5 ml tubes, followed by incubation overnight at 4 °C before storage at -80 °C.

We isolated nuclei as previously described [9] with minor modifications. Briefly, frozen tissues were placed into a 2 ml KIMBLE Dounce tissue grinder set (Sigma) with homogenization buffer consisted of 20 mM Tris pH 8.0 (Thermo Fisher Scientific), 500 mM sucrose (Sigma), 50 mM KCl (Thermo Fisher Scientific), 10 mM MgCl_2_ (Thermo Fisher Scientific), 0.1% NP-40 (Roche), 0.2U/μl RNase inhibitor (MGI), 1X protease inhibitor cocktail (Roche), 1% nuclease-free BSA, and 0.1 mM DTT. Tissues were then homogenized by 10 strokes of the loose dounce pestle. Homogenate was then strained through a 70 μm cell strainer (Falcon) and then homogenized by 10 strokes of the tight pestle to liberate nuclei. Homogenate was next strained through a 30 μm cell strainer (Sysmex) and centrifuged at 500 x g for 5 minutes at 4 °C to pellet nuclei. Nuclei were then resuspended with blocking buffer containing 1X PBS supplemented with 1% BSA and 0.2U/μl RNase inhibitor. Nuclei suspensions were then centrifuged at 500 x g for 5 minutes and resuspended with cell resuspension buffer (MGI). We counted the nuclei and made the final aliquots for snRNA-seq at a concentration of 1,000 nuclei/μl.

### Sequencing library construction using the C4 system

Single-cell RNA-seq libraries were prepared using C4 scRNA-seq kit (MGI). Barcoded mRNA capture beads, droplet generation oil and the single-cell suspension were loaded into the corresponding reservoirs on chip for droplet generation. The droplets were gently removed to the collection vial and placed at room temperature for 20 minutes. Droplets were then broken and collected by the bead filter (MGI). The supernatant was removed, and the bead pellet was resuspended with 100 μl RT mix. The mixture was then thermal cycled as follows: 42 °C for 90 minutes, 10 cycles of 50 °C for 2 minutes, 42 °C for 2 minutes. The bead pellet was then resuspended in 200 μl of exonuclease mix and incubated at 37 °C for 45 minutes. Afterward the PCR master mix was added to the beads pellet and thermal cycled as follows: 95 °C for 3 minutes, 13 cycles (for nuclei 19 cycles) of 98 °C for 20 s, 58 °C for 20 s, 72 °C for 3 minutes, and finally 72 °C for 5 minutes. Amplified cDNA was purified using 60 μl of AMPure XP beads. The cDNA was subsequently fragmented to 400-600bp with NEBNext dsDNA Fragmentase (New England Biolabs) according to the manufacturer’s protocol. Indexed sequencing libraries were constructed using the reagents in the C4 scRNA-seq kit following the steps: (1) post fragmentation size selection with AMPure XP beads; (2) end repair and A-tailing; (3) adapter ligation; (4) post ligation purification with AMPure XP beads; (5) sample index PCR and size selection with AMPure XP beads. The barcode sequencing libraries were quantified by Qubit (Invitrogen). All libraries were further prepared based on BGISEQ-500 sequencing platform [14]. The DNA nanoballs (DNBs) were loaded into the patterned nanoarrays and sequenced on the BGISEQ-500 sequencer using the following read length: 41 bp for read 1, 100 bp for read 2, and 10 bp for sample index.

### Barcode and UMI assignment, read mapping and transcript counting

For C4 scRNA-seq data, the cell barcodes (base 1 to base 10 and base 17 to base 26) and UMIs (base 32 to 41) are in read 1 and the cDNA reads are in read 2. We used the merge_fastq function of scumi [11] to extract the cell barcodes and UMI sequences from read 1, and put them in the header of their corresponding cDNA reads.

We aligned the reconstructed the FASTQ file to the reference genome using STAR [15]. For cross-species single-cell data, we used the STAR reference available in the hg19 and mm10 v2.1.0 Cell Ranger reference. For mouse cortex and lung data, we used the STAR reference available in the mm10 v1.2.0 Cell Ranger reference.

To generate a cell x gene UMI count matrix from UMI-based methods. We used the featureCounts function from the Subread packages [16] to label each alignment with a gene tag using the following parameter: -M. We then counted the UMI number for each gene in each cell using the count_umi function of scumi [11].

### Selecting the cell number

To estimate the number of cells recovered, we used the 50 percentage UMIs of the top 30th cell barcode as the threshold. The cell barcodes with UMIs greater than the threshold are considered as cells. For data generated from other platforms, the cells were selected according to the previous study [11].

### Multiplets Detection

In the cross-species experiments, the cell barcodes will be considered as the cell multiplets when UMIs from human and mouse are greater than 15%.

### Cell clustering

Seurat [17, 18] was applied to uncover the cell types in the mouse tissue datasets. For mouse lung dataset, cells with more than 200 detected genes and low than 10% mitochondrial UMIs were used for clustering. The “FindVariableGenes” function (with dispersion value higher than 0.5 and normalized expression value between 0.0125 and 3) was applied to find the highly variable genes, followed by principal component analysis (PCA) based on the highly variable genes. The top 20 PCs were used to building the *k*-NN graph by setting the number of neighbors *k* as 20, and the clusters were identified using the resolution of 0.8. The top 50 PCs were used to building the reduction map. The subpopulations of immune cells and endothelial cells were also identified using Seurat. Each cell cluster was annotated using previously reported marker genes.

For mouse cortex dataset, the data generated from other platforms were reported in Ding *et al.* [11] and were kindly provided by Dr. Joshua Z. Levin. The data matrices generated from C4 and all other platfroms were merged for downstream analysis. Cells with more than 200 detected genes and low than 10% mitochondrial UMIs were used for clustering. The “FindVariableFeatures” function (selection.methods = “vst” and nfeatures = 2000) was applied to find the highly variable genes, and the “FingIntergrationAnchors” function was applied to find the anchors in the all techniques using the top 50 PCs of canonical correlation analysis (CCA). Then integrated the different technique datasets according to the anchors, followed by principal component analysis (PCA) based on the integrated dataset. The top 12 PCs of PCA were used to building the *k*-NN graph by setting the number of neighbors *k* as 20, and the clusters were identified using the resolution of 2. The top 50 PCs of PCA were used to building the reduction map. Each cell cluster was annotated using previously reported marker genes.

## Acknowledgements

We thank Dr. Joshua Z. Levin from Broad Institute of MIT & Harvard for kindly providing raw single-cell sequencing data generated from existing platforms. This work was supported by the National Natural Science Foundation of China (No. 31900466 and No. 31900582), the Science, Technology and Innovation Commission of Shenzhen Municipality (No. JSGG20170412153009953) and the Strategic Priority Research Program of the Chinese Academy of Sciences (No. XDA16030502 and No. XDA16010114).

## Author contributions

L.L. and IJ.C. conceived the idea. C.L. and L.L. designed the experiments. IJ.C., T.W. and M.J. designed the microfluidic chip and instrument. C.L. and Z.W. designed the reagents. F.F., O.W., B.A.P. and W.W. designed and optimized the micro-particles. Y.L., L.W. and W.Wei designed the filter. L.Wu. and L.L. analyzed the data. C.L., Z.W. and Y.Y. conducted experiments. Y.Yuan., M.W., M.C., J.X., Y.Lai, P.G. and H.Z. assisted with the experiments. Q.S., X.W., S.L. and Y.Liu assisted with the data analyses. L.L. and IJ.C. wrote the manuscript. M.A.E., C.W. and G.V. revised the manuscript. X.X., Y.H., Y.Z., J.B., A.C., Z.S., M.N., W.Z. and M.A.E. provided helpful comments on this study. All authors reviewed and approved the final manuscript.

---

**Supplementary Figure 1.**
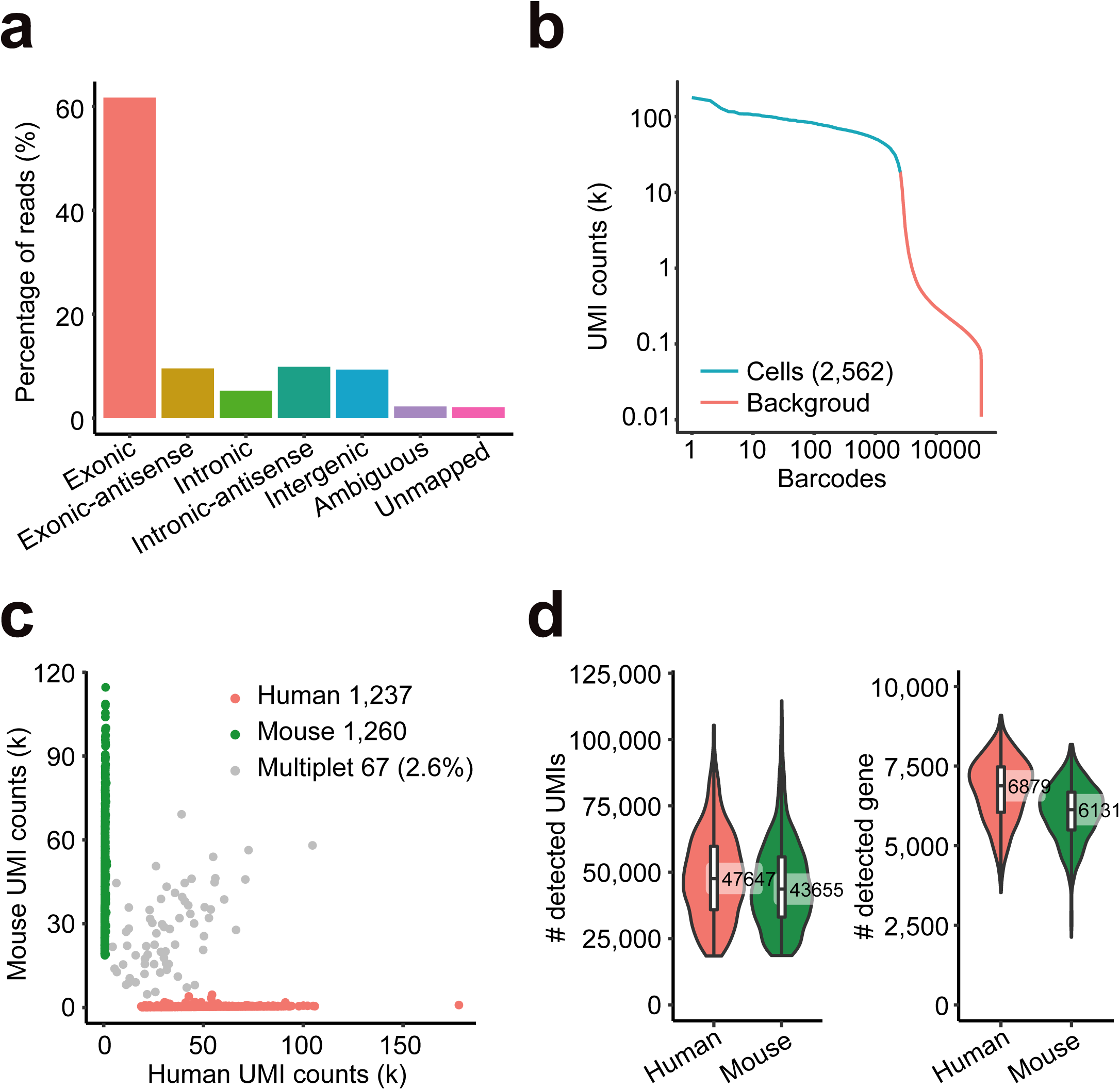
Quality metrics of C4 generated scRNA-seq. (**a**) Percentages of reads mapped to the indicated regions of the human genome. (**b**) Number of UMIs captured in each cell barcode. The cell barcodes are ordered by the UMIs number. (**c**) Scatter plot of human and mouse UMI counts detected in a mixture of 293T and NIH3T3 cells. The percentage of multiplets detected in indicated in the panel (2.6%). (**d**) Left panel: number of UMIs per cell for the human and mouse cells. Right panel: number of genes per cell for the human and mouse cells.

**Supplementary Figure 2.**
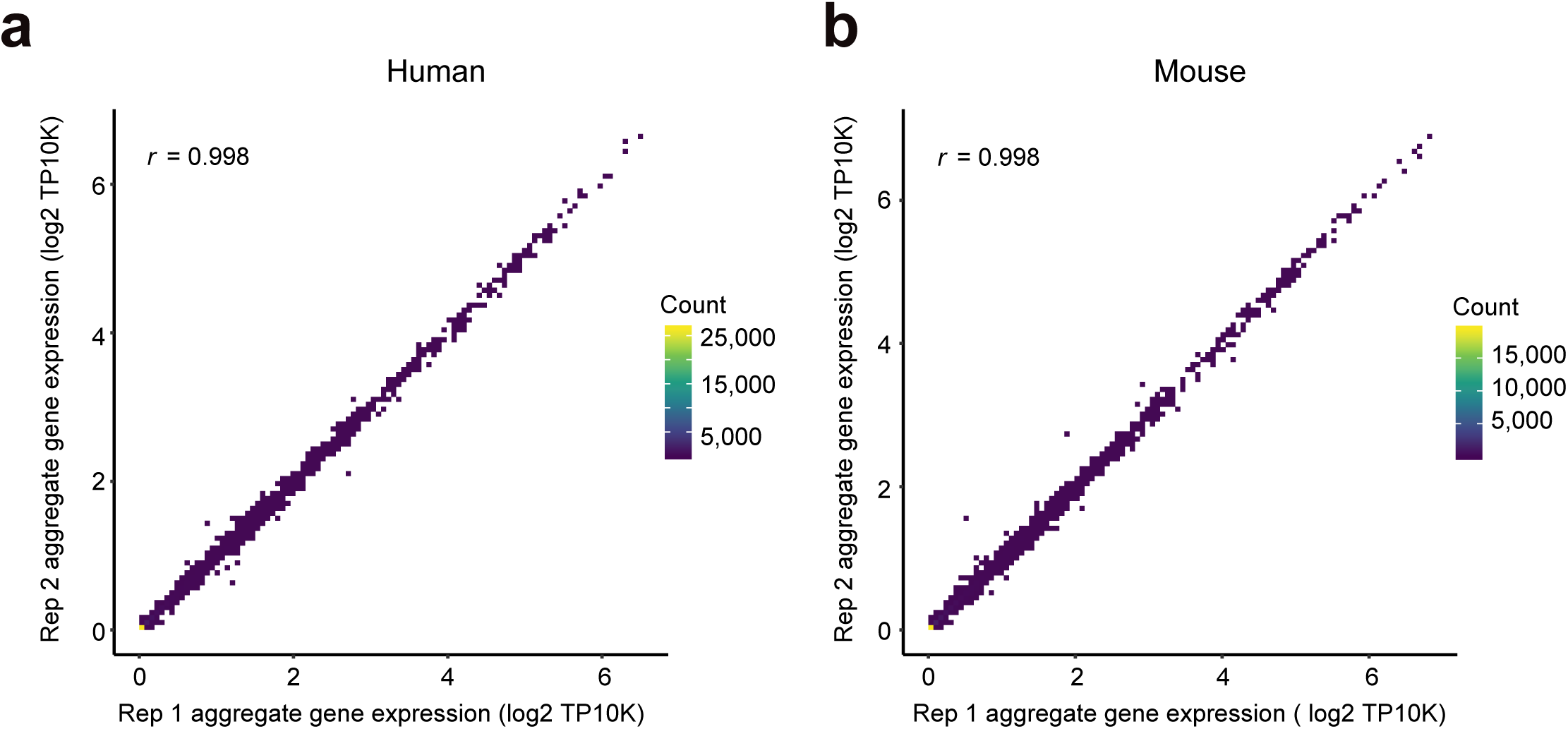
Correlation of pseudo-bulk expression profiles. The Pearson correlation coefficients between pseudo-bulk profiles of (**a**) human and (**b**) mouse cells generated from C4 system. For each replicate, we compared the log2 transcripts per 10K (log2 [TP10K+1]) for pseudo-bulk from scRNA-seq on a gene by gene basis. The color of each box in the plane is calculated from the log count of gene.

**Supplementary Figure 3.**
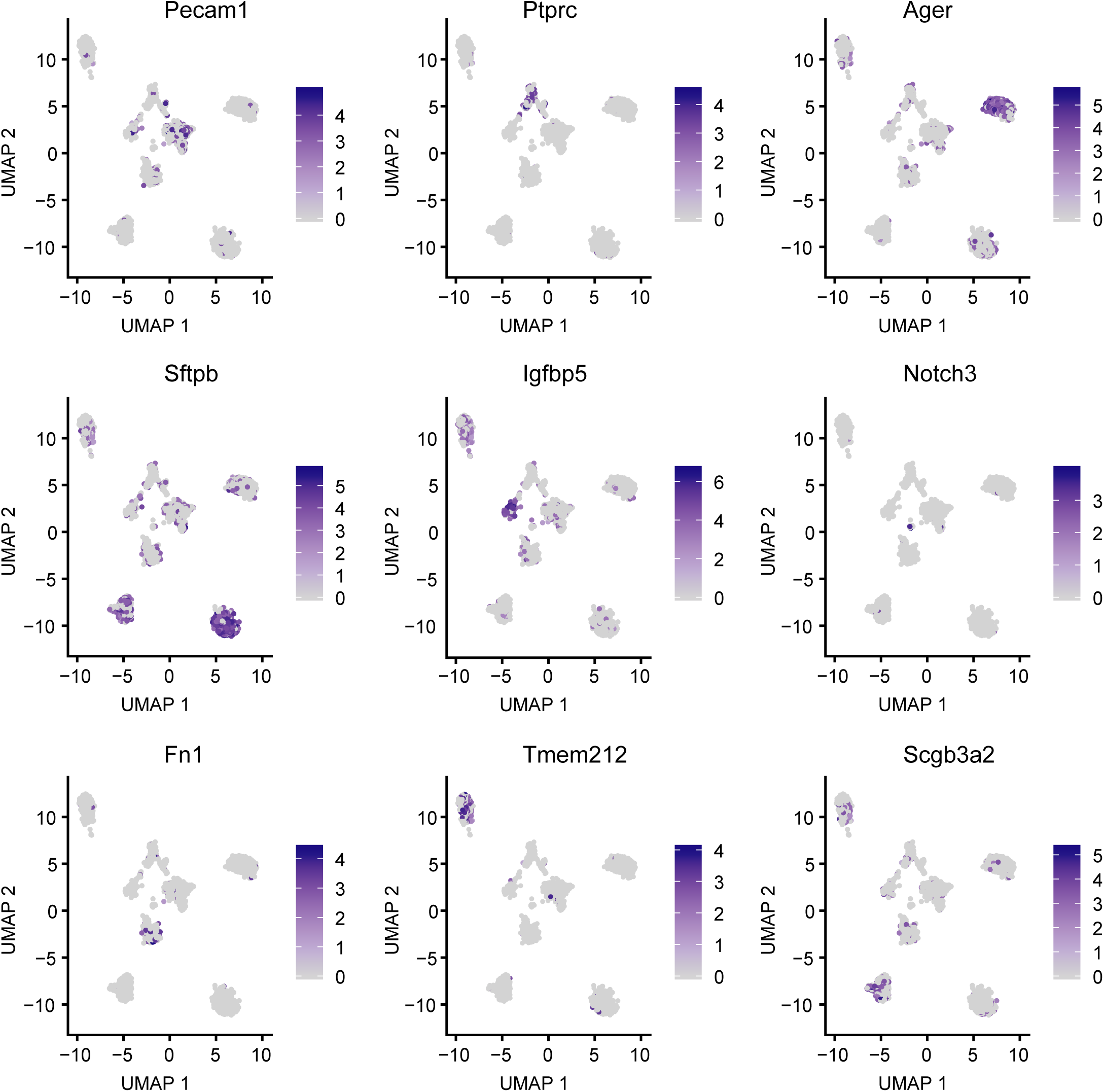
Expression patterns of the indicated marker genes in each cell type within the lung tissue. UMAP 2D scatter plot of the transcriptome of each cell. The expression value is normalized as log2 transcripts per 10K (log2 [TP10K+1]).

**Supplementary Figure 4.**
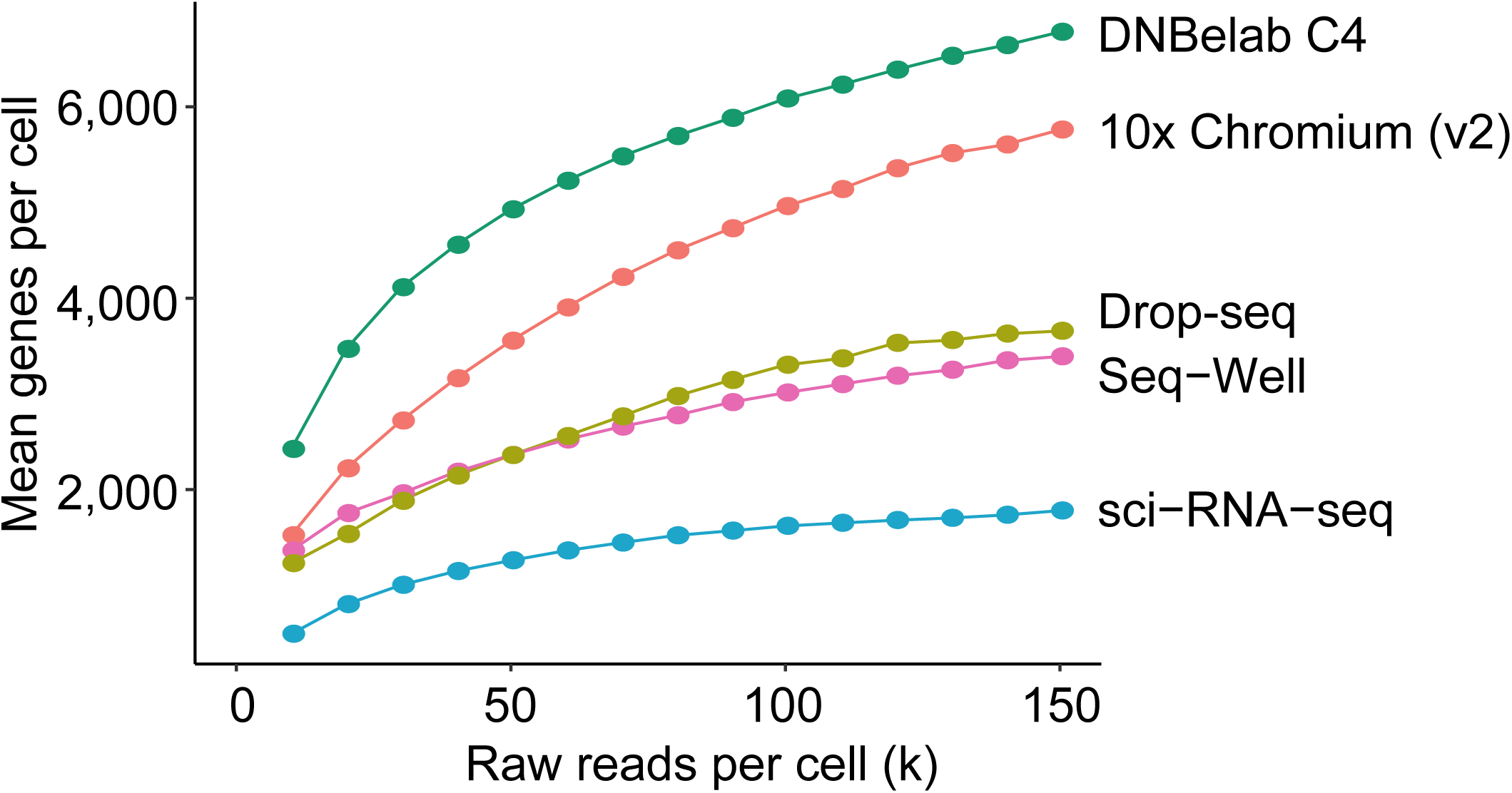
Rarefaction curves showing the effect of sequencing depth on the mean number of detected genes per cell.

